# Long live the alien: is high genetic diversity a pivotal aspect of crested porcupine (*Hystrix cristata*) long-lasting and successful invasion?

**DOI:** 10.1101/016493

**Authors:** Emiliano Trucchi, Benoit Facon, Paolo Gratton, Emiliano Mori, Nils Chr. Stenseth, Sissel Jentoft

## Abstract

Studying the evolutionary dynamics of an alien species surviving and continuing to expand after several generations can provide fundamental information on the relevant features of clearly successful invasions. Here, we tackle this task by investigating the dynamics of the genetic diversity in invasive crested porcupine (*Hystrix cristata*) populations, introduced to Italy about 1500 years ago, which are still growing in size, distribution range and ecological niche. Using genome-wide RAD markers, we describe the structure of the genetic diversity and the demographic dynamics of the *H. cristata* invasive populations and compare their genetic diversity with that of native African populations of both *H. cristata* and its sister species, *H. africaeaustralis*. First, we demonstrate that genetic diversity is lower in both the invasive Italian and the North Africa source range relative to other native populations from Sub-Saharan and South Africa. Second, we find evidence of multiple introduction events in the invasive range followed by very limited gene flow. Through coalescence-based demographic reconstructions, we also show that the bottleneck at introduction was mild and did not affect the introduced genetic diversity. Finally, we reveal that the current spatial expansion at the northern boundary of the range is following a leading-edge model characterized by a general reduction of genetic diversity towards the edge of the expanding range. We conclude that the level of genome-wide diversity of *H. cristata* invasive populations is less important in explaining its successful invasion than species-specific life-history traits or the phylogeographic history in the native source range.

## Introduction

One of the most relevant and debated questions in invasive biology concerns the importance of standing genetic diversity for successful invasions and colonization of a novel range (Reed & Frankham 2003, Frankham 2004, Facon *et al*. 2006, Roman & Darling 2007). In addition to the initial bottleneck at introduction, which may (Schmid-Hempel *et al*. 2007, Dlugosch & Parker 2008, Ciosi *et al*. 2008, Chapple *et al*. 2013) or may not (Kolbe *et al*. 2004, Roman & Darling 2007, Estoup & Guillemaud 2010, Hufbauer *et al*. 2013) decrease the genetic diversity of the introduced propagule, subsequent range expansion can also negatively affect diversity (Edmonds *et al*. 2004, White *et al*. 2013), thus likely limiting the adaptive potential of invasive populations and, ultimately, their further spread and/or persistence (Shine *et al*. 2011). Nevertheless, low neutral genetic diversity of the invasive species *per se* does not necessarily result in reduced adaptive capability (Dlugosch & Parker 2008). Rapid genetic adaptation in response to changed selective pressures encountered in the novel environment has been suggested as a possible explanation of very successful colonization events (Prentis *et al*. 2008) and, in a few cases, fast genetic changes in relevant genes have been discovered (Vandepitte *et al*. 2014). Indeed, past investigations have often reported cases of successful biological invasions despite low genetic diversity (*e.g*. Lavergne & Molofsky 2007, Hardesty *et al*. 2012). As such, other factors related to ecological traits of the exotic species and/or of the invaded ecosystem and coincidental events may be more important than initial genetic diversity in determining the success of an invasive population (Zayed *et al*. 2007). However, the vast majority of studies of biological invasions have so far utilized systems with a recent history of introduction, and thus lack a deeper temporal perspective (Strayer *et al*. 2006) leaving us with a dearth of assessments of the long-term adaptive potential of successful biological invaders (but see Cooling *et al*. 2011).

The crested porcupine, *Hystrix cristata*, was historically introduced to Italy (Italian peninsula and Sicily), making it an excellent study system to test the importance of initial genetic diversity for an invasive population to persist and spread. Ancient Romans very likely brought this animal from North Africa as an exotic pet for their villas and as a delicacy for their banquets: a genetic survey based on three mitochondrial genes identified the most likely source in North Africa (*i.e*. Tunisia) and estimated the introduction event between 2500 and 1500 years ago (Trucchi & Sbordoni 2009). Independent analyses of archaeological evidence and iconographic documentation suggested that the introduction occurred in late Antiquity or the early middle ages (1500-1200 years ago) and that the species’ presence in Sicily is not supported before early modern times (Masseti *et al*. 2010). Nevertheless, we expect a lag time between the accidental release of captive animals and their spread into the invasive range, *i.e*. when it would have been common enough to be found in archaeological sites. Based on this, we can assume that the introduction of *H. cristata* into peninsular Italy likely started around 1500 years ago, and its populations are viable and still expanding after several hundred generations. In this species, sexual maturity is achieved at the age of ca. 1 year and the following inter-litter interval is about 91-112 days (Mohr 1965, Weir 1974).

Records of the distribution of the porcupine in Italy reveal a dramatic range expansion in the last 40-50 years (Angelici & Amori 1999, Mori *et al*. 2013). During this period, the invasive population crossed the Apennines and colonized the eastern side of the Italian peninsula, passing the Po river in the Padana plain and getting as far north as the southern edge of the Alps. Intriguingly, the newly colonized area is climatically distinct from the pre-expansion range: warm temperate continental climate vs. Mediterranean coastal climate (Blasi *et al*. 2014). As the former climate type is not present in the source area (North Africa), this sudden range shift may have been driven by a novel adaptation. In stark contrast with the range expansion in North Italy, the extant populations in North Africa are currently declining due to intense anthropic pressure (Saleh & Basuony 1998; Nowak 1999; Cuzin 2003; Mohamed 2011) and the ongoing aridification of the region (Thuiller *et al*. 2006; Kröpelin *et al*. 2008). Additionally, the species still commonly occurs in Sub-Saharan Africa, from Senegal to Ethiopia and Tanzania. A sibling species, *H. africaeaustralis* (Cape porcupine), is found in austral Africa, from the Democratic Republic of Congo and Tanzania to South Africa (Nowak 1999). These two sister species are phylogenetically and ecologically very close (Mohr 1965, Trucchi & Sbordoni 2009) and their ranges of distribution largely overlap in East Africa, meaning that *H. africaeaustralis* can be used as an excellent comparison with *H. cristata* for assessing the level of genetic diversity in native populations

Employing a vast RAD sequencing dataset of more than 30,000 loci, we describe the genetic structure and diversity of the invasive Italian *H. cristata* population and compare it to the populations of *H. cristata* and *H. africaeaustralis* found in Africa. We also investigate the smaller scale genetic pattern of the Italian population that is currently expanding northwards. Finally we assess how the introduction and expansion processes have affected the genetic diversity along the whole colonization trajectory, and investigate whether high genome-wide diversity was, and still is, an important aspect of *H. cristata’s* successful invasion.

## Methods

### Sampling

A total of 280 *H. cristata* samples (244 from Italy and 36 from North-Central Africa) and 43 *H. africaeaustralis* samples from Southern Africa were collected in the fieldbetween 2004-2012 from Egypt, Ivory Coast, Ghana, Nigeria, Tanzania, Mozambique, Zambia and Namibia. Most of the samples were quills collected on the ground, but 8 of them (of *H. cristata* from Italy) were from muscle tissue (roadkill). Age and quality of preservation varied greatly across samples. DNA was extracted using the DNAse Blood and Tissue kit (Qiagen) following manufacturer's instructions. Quality and quantity of DNA was checked to identify high-quality samples suitable for genomic analysis. Concentration of DNA was measured using a fluorimetric method (Qubit, Invitrogen) and quality was checked using a spectroscopic method (Nanodrop) and by visual inspection of degradation after gel electrophoresis. Given the uneven quality and DNA preservation of the sampled materials, ca. 80% of the samples were discarded as unsuitable for genomic analyses because of strong DNA degradation, and ca. 10% of the remaining samples failed in the sequencing run.

A total of 50 samples of *H. cristata* (38 from the invasive Italian populations and 12 from native African populations) and 11 samples of *H. africaeaustralis* were selected for RAD sequencing (Fig. 1, Supplementary Table 1). Samples were grouped according to species (*H.cristata:* H.cri; *H. africaeaustralis:* H.afr) and geographical origin (H.cri-SS: Sub-Saharan Africa; H.cri-NA: North Africa; H.cri-IT-Sicily: Sicily, Italy; H.cri-IT-south: South Italy; H.cri-IT-centre: Central Italy; H.cri-IT-north: North Italy). In addition, samples in H.cri-IT-north were further grouped according to the year of establishment of a population (Angelici *et al*. 2003, Mori *et al*. 2013) in the area they were sampled (H.cri-IT-north-1959: distribution in 1959; H.cri-IT-north-1999: distribution in 1999; H.cri-IT-north-2012: distribution in 2012).

**Figure 1.**
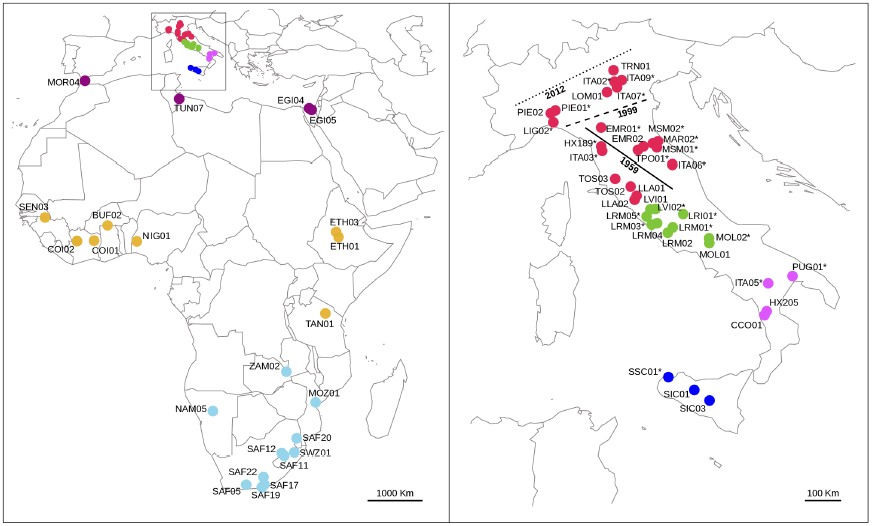
Distribution of the genotyped individuals in Africa (left panel) and Italy (right panel). Colors identify *a priori* groups defined in the text: H.afr (light blue), H.cri-SS (orange), H.cri-NA (purple), H.cri-IT-Sicily (blue), H.cri-IT-south (light purple), H.cri-IT-centre (green), H.cri-IT-north (red). The recent range expansion documented in North Italy is also shown: the black solid line represents the north-easternmost distribution in 1959 (H.cri-IT-north-1959); the large-dashed black line represents the northernmost distribution in 1999 (H.cri-IT-north-1999) and the small-dashed black line represents the northernmost distribution in 2012 (H.cri-IT-north-2012) according to Angelici & Amori (1999) and Mori *et al*. (2013). Individuals from Italy included in both the *global* and the *invasive_reduced* dataset are not marked with an asterisk; in the text we refer to this group of samples as H.cri-IT.

### RAD sequencing

The RAD sequencing protocol from Baird *et al*. (2008) was slightly modified to prepare the libraries. Approximately 100 ng of genomic DNA per sample were digested with the restriction enzyme *Sbf*I (NEB). Each sample was ligated to a unique barcoded P1 adapter prior to pooling in a single library. Libraries were sheared by sonication on a Bioruptor (Diagenode) where the target size range fraction of 300-500 bp was achieved after seven cycles of sonication (30 seconds ON, 30 seconds OFF). After concentration to 25 μl by DNA capture on magnetic beads (beads solution: DNA = 0.8: 1), libraries were size selected by gel electrophoresis and manual excision. Capture on magnetic beads (beads solution: DNA = 0.8: 1) was then employed in all following purification steps (*i.e*. after blunt-end repairing, poly-A tailing, P2 adapter ligation and library enrichment by PCR). To reduce amplification bias, PCR reactions were split in 8 x 12.5 μl aliquots per library, separately amplified and then pooled again. Libraries were quantified by a fluorimetric-based method (Qubit, Invitrogen) and molarity was checked on an Agilent Bioanalyzer chip (Invitrogen). A final volume of ca. 20 μl per library with a DNA concentration of 20-25 ng/μl was submitted for paired-end 100 bp sequencing (two lanes on a ILLUMINA HiSeq2000) at the Norwegian Sequencing Centre, University of Oslo.

### Bioinformatic analyses

Raw reads were processed using the scripts included in the *Stacks* package (Catchen *et al*. 2013) on the ABEL cluster server facility at the University of Oslo. Raw reads were quality filtered and demultiplexed according to individual barcodes using the script *process_radtags.pl* in the *Stacks* package with default settings. Cleaned reads were then aligned into loci and SNPs called across individuals using the script *denovo_map.pl* in the *Stacks* package: the minimum coverage to call a stack of identical reads was set to 10 (option-m), the maximum number of mismatches allowed when joining stacks into the same locus to 7 (option-n) and the maximum number of mismatches allowed when joining loci across individuals to 7 (option-N). In order to capture different levels of genetic variability within and between populations of the two species, we built three separate catalogs: *i*) one catalog including all Italian and North Africa individuals (*invasive* dataset, 42 individuals), *ii*) one catalog including a subset of the Italian individuals taking into account both an even representation of the invasive range and the highest average coverage across genotyped loci (*invasive_reduced* dataset, 16 samples: H.cri-IT; Fig. 1), and *iii*) one catalog including the same subset of high-coverage Italian individuals as in the previous catalog as well as all of the African individuals (*global* dataset, 39 individuals). We included only a small subset of the invasive samples (16) in the *global* catalog in order to avoid overweighting the genetic diversity in the invasive population, where the largest proportion of samples were analyzed, and to minimize the SNPs ascertainment bias in this dataset as a result. The function *export_sql.pl* in the *Stacks* package was used to extract loci information from each catalog applying a global threshold of 25% to the maximum number of missing sample per locus and 10 to the maximum number of SNPs per locus. This high threshold for the maximum number of SNPs per locus, especially in the case of the invasive population, was justified by the consideration that filtering loci on the basis of their shallow variation might introduce a severe bias towards recent coalescent events. In the case of the invasive population, the only possibility to retrieve the genetic signature of a bottleneck is to find divergent alleles whose coalescence time is deeper than the bottleneck itself and that can provide information about the pre-bottleneck ancestral population size. In addition, the accumulation of substitutions along the genome is a random process with a mean rate and extreme cases are expected. A careful screening of putative paralogous loci or non-random distribution of SNPs along each locus was then performed as follows. We wrote custom python scripts (see Supplementary Materials) to further filter the dataset in order to exclude loci with more than 2 alleles per individual, with heterozygosity above 0.75, or deleveraged by the Stacks algorithm. We found an increase in the number of SNPs called in the last 10 positions across all of the loci (Fig. S1); trimming the raw reads of 10bp and performing again the SNP calling produced the same pattern. We believe this is an artifact of the SNP calling algorithm, so any SNP recorded in the last 10 base pairs of each locus was considered unreliable and discarded. Further filtering to reduce missing data was applied on a case-by-case basis, according to each downstream analysis.

### Genetic structure of native and invasive populations

The *global* dataset was used to infer the overall structure among native and invasive populations of the two porcupine species. Following Wagner *et al*. (2013), loci were concatenated into a single sequence per sample, coding heterozygous sites as ambiguities in agreement with the IUPAC code. The whole sequence of each locus was included in order to get empirical estimates of base composition and percentage of invariant sites. A Maximum-Likelihood algorithm with a GTR * G * I substitution model was employed to reconstruct the phylogenetic tree of our samples using 100 rapid bootstrap inferences and thereafter a thorough ML search in *RAxML* 7.2.8 (Stamatakis 2006). Results were visualized and edited in *FigTree* 1.4 (http://tree.bio.ed.ac.uk/software/figtree/). Although this cannot be considered a true phylogenetic reconstruction, this random concatenation of recombining genomic fragments has proven to be informative (Wagner *et al*. 2013).

The *invasive* dataset was then used to study the fine-scale geographic structure of the *H. cristata* invasive population and its relationship with North Africa source population. First, using all of the SNPs in each locus and all of the samples in the dataset (H.cri-IT and H.cri-NA; Fig. 1), we performed a Neighbor-Joining Network (NeighborNet) analysis (Bryant & Moulton 2004) based on uncorrelated *p*-distances in *Splitstree* (Huson & Bryant 2006). We then performed a Principal Component Analysis (PCA) reducing our dataset to only one randomly-chosen SNP per locus. To better capture the variance in the invasive range, only individuals belonging to the invasive population (H.cri-IT; Fig. 1) were included in the PCA, while individuals from North Africa were excluded. Given that PCA is very sensitive to missing data (in this case, missing loci in each individual), we further filtered our dataset, removing four individuals with more than 50% missing loci (EMR01, ITA09, SSC01, LRM03). The strict filtering in this second analysis was necessary because samples with too many missing loci tend to be unassigned (e.g. appear at the centre of the axes). The *glPca* function in the *R* packages *“adegenet”* was used for calculations.

### Demography of the invasive population

We used RAD loci genotyped in the high-coverage Italian individuals (H.cri-IT; Fig.1) to reconstruct the demographic history of the invasive population following an approach recently proposed by Trucchi *et al*. (2014) that has been proven to be particularly efficient in reconstructing recent demographic events (Tutorial available at http://www.emilianotrucchi.it/images/EBSP_RADseq_tutorial.pdf). In short, a selection of highly variable RAD loci with more than 3 SNPs per locus are used as short sequences in a coalescent-theory based multi-locus analysis (Extended Bayesian Skyline plot; Heled & Drummond 2008) implemented in *BEAST* 1.7.4 (Drummond & Rambaut 2007). Four random selections of 50 loci with 4 to 9 SNPs (no loci with more than 9 SNPs passed our filtering criteria) from the *invasive_reduced* dataset were used in four replicated runs and checked for convergence. Analyses were performed as follows: *i*) nucleotide substitution models, clock models and tree prior models were unlinked across loci; *ii*) the nucleotide substitution model was set as a HKY with empirical base frequency; *iii*) a strict molecular clock was set for each marker with a uniform prior distribution of the substitution rate bounded within 0.5 and 0.005 sub/s/Myr; *iv*) the Extended Bayesian Skyline Plot (EBSP) was selected as a tree prior model. 200 million iterations were set as run length. In addition, we ran the EBSP analyses adding the mtDNA Control Region sequences of the invasive population published in Trucchi & Sbordoni (2009) to a random selection of 50 RAD loci. We used this analysis to calibrate our demographic reconstruction, applying a substitution rate of 0.2 substitution/site/Myr to the mtDNA marker only and leaving the substitution rates for the RAD loci to be estimated in the analysis. The substitution rate employed for the calibration was derived from the rate estimated in another rodent species (Mus *musculus;* Rajabi-Maham *et al*. 2008) and corrected for the longer generation time in porcupines. An HKI with Gamma (4 classes) and Invariant site substitution model and a strict clock model were implemented for the mtDNA Control Region. Three replicates were checked for convergence. All analyses were run on a 24 CPUs server at the University of Oslo. Results were checked on *Tracer* 1.6 (http://tree.bio.ed.ac.uk/software/tracer) and the plot of the EBSP data was drawn in *R* (R Development Core Team 2011). To take into account the structure in the invasive population, we also repeated the EBSP analysis using samples from only one of the groups (H.cri-IT-north).

### Heterozygosity estimates

Events of introduction into a new range are expected to induce a reduction in heterozygosity in the invasive population, and quick range expansions in the invasive range could produce a further clinal reduction in heterozygosity towards the expansion front. Nevertheless, levels of individual heterozygosity ultimately depend on levels of inbreeding (Hoffman *et al*. 2014), and are influenced by a variety of demographic process at different temporal and spatial scales. Observed and expected heterozygosity were estimated in native and invasive populations/groups of individuals (Fig.1) using the *global* and the *invasive* datasets. To test for the effects of the introduction event, we used the *global* dataset, using the following partition of native and invasive individuals: H.afr, H.cri-SS, H.cri-NA, and H.cri-IT. To test for local effects in the different invasive populations, we used the *invasive* dataset and the groups H.cri-IT-north, H.cri-IT-central, H.cri-IT-south, H.cri-IT-Sicily, and H.cri-NA. To test for the effects of the recent range expansion in north Italy, we used the *invasive* dataset and the groups H.cri-IT-north-1959, H.cri-IT-north-1999, and H.cri-IT-north-2012.

Despite their relevance as proxies for levels of inbreeding (Slate & Pemberton 2002), accurate estimates of individual observed heterozygosity from multi-locus data are difficult to calculate (see Aparicio *et al*. 2006 for a review). We first estimated individual observed heterozygosity as the number of heterozygous loci divided by the number of genotyped loci in each individual (H_o ind_). As our RADseq loci were not mapped to a reference genome, a fraction of non-orthologous loci could have been included in the analysis and may appear as loci with artificially high heterozygosity. To address this issue, we down-weighted the contribution of each locus according to its observed heterozygosity across individuals (H_o ind-norm_). To apply this correction, we first calculated the observed locus-by-locus heterozygosity across all of the individuals in the dataset (H_o loc_ = number of heterozygous genotypes / number of genotype at a locus). Then we re-calculated the individual observed heterozygosity correcting the contribution of each locus by a factor 1 - H_o loc_. Another important aspect to take into account when estimating observed individual heterozygosity from next-generation data is that the probability of scoring a heterozygous genotype at a locus is a function of its depth of coverage. To correct for this, we included the coverage depth of each individual as a predictor of individual observed heterozygosity in a linear model and we calculated the deviation of each individual from the model (*i.e*., analysis of model’s residuals). Individual-based residuals were then grouped as described above and compared. An additional analysis using log-transformed values for depth of coverage was also performed to take into account the fact that the probability of retrieving both alleles at a heterozygous locus is expected to reach a plateau. The significance of differences in observed heterozygosity among geographical groups of samples was assessed by F-test comparing a linear model where observed heterozygosity was predicted by both the number of reads and the geographic origin of the sample with a reduced model with the number of reads as the only predictor. When the F-test revealed at least a statistical trend (P < 0.1), the statistical significance of all pairwise contrasts among geographically-defined groups of samples was calculated by running the full linear model N-1 times (with N = number of groups) and setting, each time, a different group as the reference level for the 'group' factor. Analyses were performed using custom python scripts (available as Supplementary Material) and basic *R* functions.

Expected heterozygosity (*i.e*., gene diversity; He) was also estimated for each of the groups described above according to the formula in Nei (1978). A minimum number of 3 individuals genotyped per group was required to include a locus in the analysis. As compared to H_o-ind_, H_e_ is less biased by the individual sequencing coverage because rare alleles in the population still have 50% probability to be sequenced in those individuals that carry them. Even if information on the population structure in the native range was scarce, estimation of population-level Watterson's theta and π were also calculated for both native and invasive populations. Analyses were performed using custom python scripts (available as Supplementary Material).

## Results

### RAD sequencing data

After de-multiplexing, the average number of single reads retained per individual was between ca. 500,000 and ca. 7,100,000. The starting quality of the DNA strongly influenced this inequality across individuals, with extractions from muscle tissue resulting in a much higher sequencing yield than extractions from quills. The total number of loci retained in each catalog after filtering was: 17,504 in the *invasive_reduced*, 30,506 in the *invasive* and 19,559 in the *global* dataset with an average coverage per allele per individual of ca. 15X. Distribution of SNPs per position and of individuals genotyped per locus is shown in Figure S2.

### Genetic structure of native and invasive populations

The Maximum-Likelihood tree reconstructed using the information provided by the *global* dataset showed a clear differentiation between *H. africaeaustralis* and *H. cristata* (Fig. 2); samples belonging to *H. africaeaustralis* formed a cluster of mostly unresolved relationships with little or no geographical structure in the genetic diversity. On the other hand, *H. cristata* populations showed a marked geographic differentiation. In this species, the genetic diversity mirrored its distribution from East Africa (Tanzania and Ethiopia), where *H. cristata* overlaps with *H. africaeaustralis*, to the West (Nigeria, Ivory Coast, Burkina Faso and Senegal), and then northward, to the Mediterranean coast of Africa and Italy (Morocco, Tunisia, Egypt, Italy). In the invasive range, the genetic structure of *H. cristata* samples clearly followed their geographic origin with clusters of individuals corresponding to H.cri-IT-north, H.cri-IT-centre, H.cri-IT-south, and H.cri-IT-Sicily. The latter also included individuals from Tunisia and Egypt (from the group H.cri-NA).

**Figure 2.**
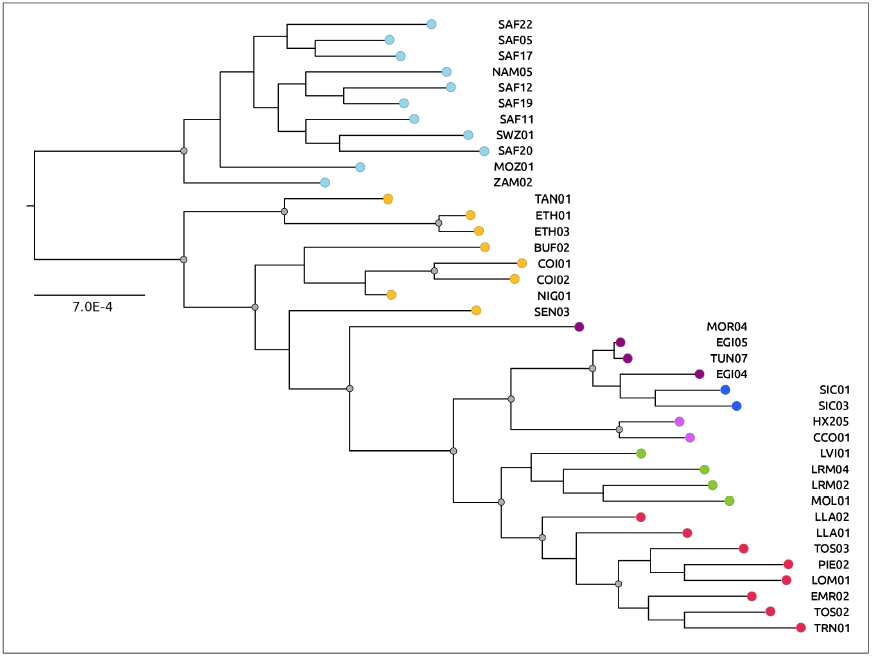
Global genetic structure. Maximum-Likelihood tree built using the *global* dataset (all individuals not marked with an asterisk in Africa and Italy in Fig.1). Nodes with bootstrap support of > 99% are shown (grey filled circles). Colors represent the same geographic locations as in Fig. 1.

The structure of the genetic diversity in the *H. cristata* invasive range was investigated in more detail using the *invasive* dataset. Results of the Neighbor-Joining Network analysis (Fig. 3a) and of the PCA (Fig. 3b) were consistent with each other and with the Maximum-Likelihood tree estimated on the *global* dataset. The same four clusters, as described above, clearly characterized the invasive range. Only two individuals did not cluster according to their geographic origin (Fig.1): ITA09 (from the group H.cri-IT-north), sampled at the early stage of the range expansion in the northernmost area occupied in Italy, and MOL02 (from the group H.cri-IT-centre), sampled in central Italy. In all of the analyses, these two samples clustered in H.cri-IT-centre and H.cri-IT-north, respectively.

**Figure 3.**
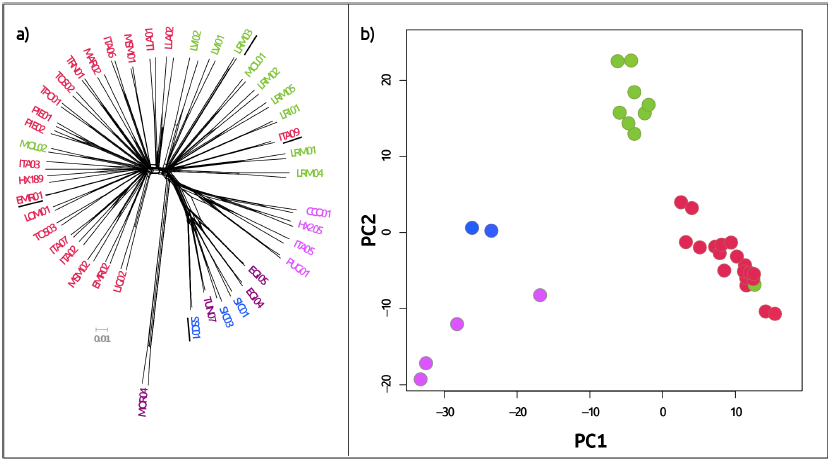
Structure and diversity in the invasive range. Neighbor-Joining Network (a) and principal component analysis (b) using the *invasive* dataset (See Methods for details). Some individuals (EMR01, ITA09, SSC01, LRM03; underlined in panel a) are included only in the Neighbor-Joining Network but not in the PCA; they were discarded according to the strict filtering required for this analysis. Colors represent the same geographic locations as in Fig. 1.

### Demography of the invasive population

The reconstruction of the past demography of the invasive population showed a clear signature of a recent, although mild, bottleneck (Fig. 4). According to our result, the coalescent effective population size at introduction corresponded to few hundred individuals, and it was only 5 times lower than before. The time of the bottleneck is estimated to be within the last 2000 years when taking into account the 95% confidence intervals. The calibrated demography showed a decrease in population size starting well before the putative introduction event, this decrease corresponds to the end of the African Humid Period (AHP), when the green Sahara was turning into a desert (deMenocal *et al.*, 2000). The bottleneck was also found when the RAD loci only and no mtDNA markers were included in the analyses (Fig. S3), but it was not detectable when the mtDNA was analyzed on its own (Trucchi & Sbordoni 2009). Independent runs employing only individuals from H.cri-IT-north produced the same demographic reconstruction, ruling out structure in the invasive population as a source of the observed pattern (not shown).

**Figure 4.**
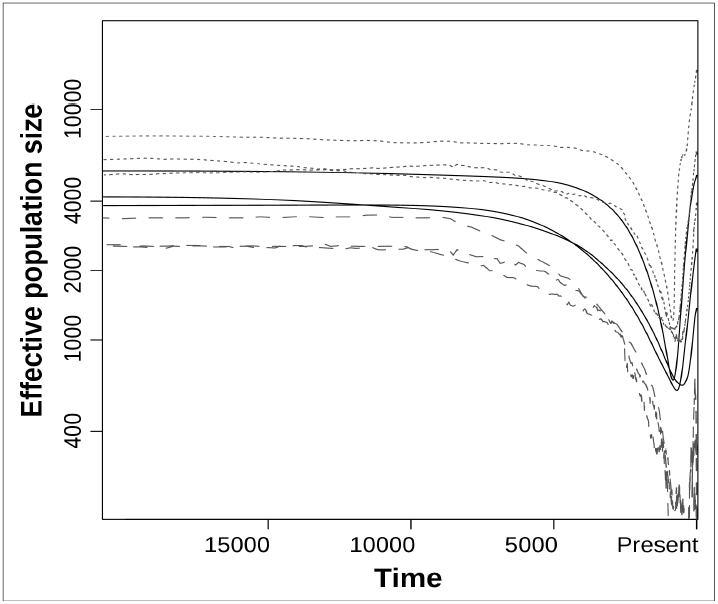
Recent demography of the invasive population. Extended Bayesian Skyline plot using 16 selected individuals from the invasive range not marked with an asterisk in Fig.1 (*invasive_reduced* dataset; see Methods for details). Results of three independent runs including 50 RAD loci and one mitochondrial locus are shown. The median (black), as well as the lower (gray large-dashed) and the upper (gray small-dashed) boundaries of the 95% credible region are shown. Time is shown in years before present (x-axis) while effective population size is in individuals (y-axis).

### Heterozygosity estimates

Estimates of individual observed and population expected heterozygosity are shown in Figure 5. Robustness of this analysis to the amount of missing data, to the correction for highly-heterozygous loci (H_o ind-norm_), and to the log-transformation of the predictor (log-reads) is reported in Figure S4-S5. Considering both observed and expected estimates, heterozygosity was significantly higher (p < 0.001 considering all comparisons) in H.afr and H.cri-SS than in H.cri-NA and H.cri-IT (Fig. 5a). In the invasive range, the level of H_o ind_ was similar across all of the groups considered in the analysis (p = 0.248 considering all comparisons), only slightly lower in H.cri-IT-south and H.cri-IT-Sicily (Fig. 5b). When the predictor is not log-transformed (p = 0.137 considering all comparisons), H_o ind_ in H.cri-NA and H.cri-IT-Sicily was slightly lower and higher than before, respectively (Fig. S5). However, it has to be noted that both groups included only a few individuals (four and three respectively). In contrast to H_o ind_, He was lower in H.cri-IT-south and H.cri-IT-Sicily than in the other groups. When investigating the fate of the heterozygosity along the recent range expansion in H.cri-IT-north (Fig. 5c), our analysis showed a consistent decrease in both observed and expected heterozygosity from the core to the edge of the recently colonized area (p = 0.058 considering all comparisons). The decrease was particularly evident in the most recently occupied area (H.cri-IT-north-2012). Mutation-scale effective population size (theta) in the native and invasive range was estimated using the two alternative estimators π and Watterson's: 0.27 and 0.30 in H.afr; 0.35 and 0.32 in H.cri-SS; 0.22 and 0.18 in H.cri-NA; 0.22 and 0.20 in H.cri-IT.

**Figure 5.**
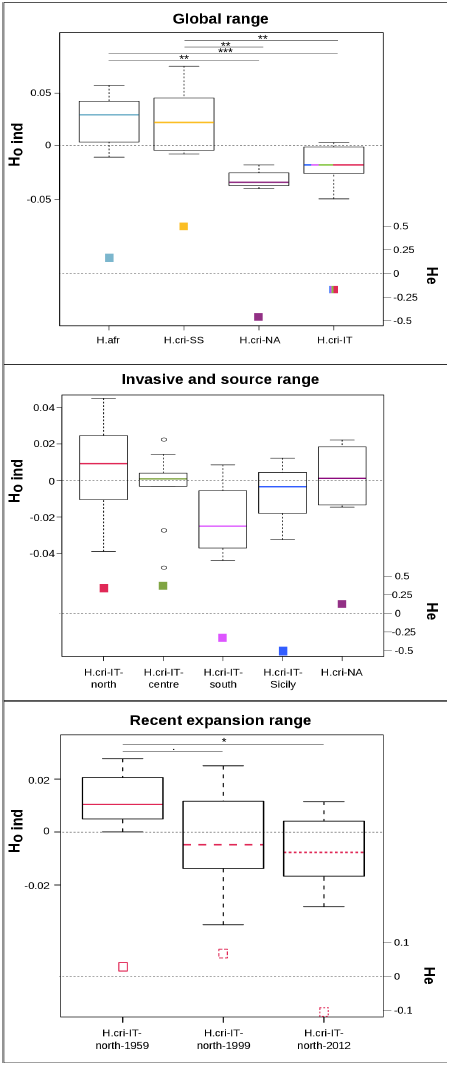
Observed and expected heterozygosity in native and invasive populations. Residuals of the model H_o ind_ ~ log(raw reads) and H_e_ as deviation from the mean are analyzed across the native and invasive range (p < 0.001 considering all comparisons) using the *global* dataset (a), across the invasive and the source native range (p = 0.237 considering all comparisons) using the *invasive* dataset (b), and across the recent expansion range (p = 0.058 considering all comparisons) using the *invasive* dataset (c). Pairwise significance between populations/groups is reported only in case the p-value for the geographic factor across all comparisons was below 0.1 (a and c panels). P-value pairwise: '.' < 0.1; '*' < 0.05; '**' < 0.01; '***' < 0.001. Colors and patterns represent the same geographic locations as in Fig. 1.

## Discussion

### Pivotai drivers of successful invasions

The *H. cristata* populations introduced in Italy more than 1500 years ago - and still expanding - are the result of an extremely successful invasion (Trucchi & Sbordoni 2009; Masseti *et al*. 2010; Mori *et al*. 2013). Our analyses demonstrate that both genetic diversity, measured as expected heterozygosity (H_e_), and inbreeding, measured as individual observed heterozygosity (H_o ind_), in the invasive populations are lower than in the natural populations of both *H. cristata* and *H. africaeaustralis* from Sub-Saharan and south Africa (Fig. 5a). However, genetic diversity in the native populations of *H. cristata* from the source range in North Africa is similar or slightly lower than in the invasive range. Including only four individuals from different geographic areas, the sample from North Africa is rather small and does not allow us to get conclusive information on the genetic diversity of the source population. Nevertheless, it has to be noted that individuals from Sicily, clustering together with those from Tunisia and Egypt (Fig. 3), show very similar levels of genetic diversity (Fig. 5b) and support the hypothesis that genetic diversity in the source population was lower than in other Sub-Saharan populations. On the other hand, the main finding of our study (i.e. successful invasion and colonization despite low genetic diversity) is supported by the comparison between the invasive population and the two native populations from Sub-Saharan and south Africa. Relative effective population sizes of native and invasive populations, calculated both on the basis of segregating sites and pairwise differences, are consistent with the heterozygosity estimates and also support the conclusion that the introduction bottleneck left a minor, if not negligible, signature on the invasive population. The effective population size in the source population (H.cri-NA) is lower as compared to other native populations (H.cri-SS and H.afr) supporting a past history of contraction and fragmentation in the North Africa range.

Our demographic reconstruction of the invasive populations clearly showed a mild bottleneck (with a few hundred individuals at the lowest point) that can be calibrated at the introduction time (Fig. 4). Nevertheless, the marked genetic structure found in the invasive range (Fig. 3) supports a multiple introduction scenario (in contrast to a single massive introduction and naturalization of hundreds of individuals), where many small propagules were introduced over time by continuous commercial trading instead of. Indeed, Sicily and South Italy could have even been colonized at a later stage as suggested by the higher genetic (Fig. 2, 4) and morphological (Angelici *et al.*, 2003) similarity with North African populations and further supported by the analysis of historical records (Masseti *et al*. 2010). Interestingly, very limited admixture among different propagules should have characterized the demographic dynamics of the invasive populations in order to retrieve such clear structure today. Thus, increased genetic diversity through admixture in the newly colonized area was not necessary for the long persistence of this invasive species. In such cases, limiting connectivity among different invasive propagules would be of little to no help in preventing the colonization (see Rius and Darling 2014 for a review).

Another important implication of our study is that initial genome-wide diversity does not necessarily explain an invasive species’ success. Even if high genetic diversity appears to be positively correlated with success in the early phases of colonization (Forsman 2014), the correlation between genetic diversity and long-term viability (and evolvability) of an invasive species could be less straightforward than it intuitively seems (Dlugosch *et al*. 2015). Indeed, several studies described successful invasions despite an initial low genetic diversity (*e.g*. Zayed *et al*. 2007, Harrison & Mondor 2011) or, alternatively, reporting an increased diversity in the invasive range (*e.g*. Lavergne & Molofsky 2007, Signorile *et al*. 2014). We suggest that a thorough understanding of the phylogeographic history as well as the phenotypic and genomic traits of the source populations is necessary to clearly identify common features of highly invasive species.

In the case of *H. cristata*, introduced individuals were likely sourced from North African populations that already had low genetic diversity. In fact, North African populations were (and still are) suffering from the ongoing desertification of the Sahara, which started approximately 6000 years ago (deMenocal *et al*. 2000). This long history of habitat fragmentation after the African Humid period is likely to have caused contraction and isolation of North African populations, thus increasing inbreeding and decreasing (local) genetic diversity. One of the short-term effects of inbreeding is an increased probability that an individual could carry recessive deleterious alleles in homozygosity, decreasing its fitness and, in general, reducing the viability of the small population (Charlesworth & Willis 2009). Nevertheless, if a small population survives a long period of inbreeding, the expectation is that some (or most) of the deleterious alleles have already been exposed to selection and likely purged from its gene pool (Crow 1970), making that population more resistant to further inbreeding (but see Crnokrak & Barret 2002 for a review). This has been shown in laboratory experiments where higher fitness was found in invasive populations that experienced mild bottlenecks and high inbreeding in the past compared to native populations that never experienced a bottleneck (Facon *et al*. 2011, Tayeh *et al*. 2013). The long history of fragmentation and isolation of *H. cristata* in North Africa could have then favoured the purging of several deleterious alleles in the source gene pool. Individuals originating from these populations could have been less susceptible to the negative effects of small population size at introduction and, as a result, more efficient in establishing a viable population in the invasive range.

### Recent invasive evolutionary dynamics

In this study, we further demonstrate that the dramatic range expansion of *H. cristata* recorded in the last 50 years in Italy is following a leading-edge pattern (Hewitt 1996), with the north population acting as the only colonization source (Fig. 1, 2, 4). One exception in our dataset is the sample collected in North Italy (ITA09), at an early stage of the range expansion, which is genetically similar to the Central Italy population (H.cri-IT-centre). This mismatch could be explained by labeling error, or could actually be the result of long-distance dispersal due to human-mediated translocation of individuals (Mori *et al*. 2013). The same reasoning could be applied to the sample in Central Italy (MOL02) that clusters with the northern group.

Our analyses reveal a clear decrease in the genetic diversity with the year of colonization (Fig. 5c), mainly as individual observed heterozygosity but also in terms of expected heterozygosity at the northernmost edge of the expansion. Nevertheless, the overall pattern of genetic diversity is consistent with expectations drawn from the gene surfing model (Edmonds *et al*. 2004, DeGiorgio *et al*. 2011). According to this model, genetic diversity is reduced at the leading edge of the expansion, whereas local gene flow and admixture is expected to balance the diversity loss at the trailing edge. However, if there is any spatial constraint in the newly colonized area (mountain ridges, rivers), it will be more difficult to re-establish the level of diversity present at the original core of the expansion (Excoffier *et al*. 2009). The mountain ridge along the Italian peninsula (Apennines) and the main river in the Padana plain (Po) could act as constraints to future gene flow in the expansion range. It remains to be investigated what the main driver of the observed range expansion is. At least three hypotheses can be proposed as an explanation: *i*) the reduced anthropic pressure due to the wide-scale abandonment of the countryside after the Second World War and the legal protection of this species since 1980; *ii*) the effect of ongoing climate change (for analyses of climate change in Italy cf. Brunetti *et al*. 2006); *iii*) an emerging adaptation in the north population that allowed the colonization despite relatively novel climate conditions (from Mediterranean sub-coastal to warm temperate areas; Blasi *et al*. 2014). Based on the solid background about the structure and the dynamics of the neutral genetic diversity provided in this study, further analyses should focus on putative adaptive response(s) in the expanding invasive population, aiming at disentangling the effects of demographic processes from those of selection.

## Acknowledgements

We would like to thank all of the colleagues and friends who kindly helped providing samples from all over Africa. We thank Anna Mazzarella for helpful comments on the early version of the manuscript and four anonymous reviewers for their excellent contribution. This study was supported by Marie Curie Intra European Fellowships (FP7-PEOPLE-IEF-2010, European Commission; project no. 252252 to E.T.) and by the Centre for Ecological and Evolutionary Synthesis, Department of Biosciences, University of Oslo, Norway.

## Data accessibility

Raw ILLUMINA reads for each sample have been uploaded to the SRA (Acc. num. SRP065809). Processed RAD datasets are publicly available on Dryad repository: doiXXXX

## Author contributions

Designed research: ET, NCS; performed research: ET; contributed analytical tools: ET, PG; contributed sampling ET, EM, PG; analyzed data: ET, PG; wrote the paper: ET, BF, PG, NCS, SJ.

